# In Search of Consciousness: Examining the Temporal Dynamics of Conscious Visual Perception using MEG time-series data

**DOI:** 10.1101/603043

**Authors:** Anh-Thu Mai, Tijl Grootswagers, Thomas A. Carlson

## Abstract

The mere presence of information in the brain does not always mean that this information is available to consciousness (de-Wit, Alexander, Ekroll, & Wagemans, 2016). Experiments using paradigms such as binocular rivalry, visual masking, and the attentional blink have shown that visual information can be processed and represented by the visual system without reaching consciousness. Using multivariate pattern analysis (MVPA) and magneto-encephalography (MEG), we investigated the temporal dynamics of information processing for unconscious and conscious stimuli. We decoded stimulus information from the brain recordings while manipulating visual consciousness by presenting stimuli at threshold contrast in a backward masking paradigm. Participants’ consciousness was measured using both a forced-choice categorisation task and self-report. We show that brain activity during both conscious and non-conscious trials contained stimulus information, and that this information was enhanced in conscious trials. Overall, our results indicate that visual consciousness is characterised by enhanced neural activity representing the visual stimulus, and that this effect arises as early as 180 ms post-stimulus onset.

## 1. Introduction

The human visual system processes a steady stream of inputs, but only a subset of this information enters consciousness. This dissociation between perceptual processing and visual consciousness has been studied extensively using paradigms such as masking (Breitmeyer & Öğmen, 2006), and binocular rivalry (Blake, 1998). In these studies, consciousness denotes visual awareness of a stimulus in the environment, which differs from the physiological state of wakefulness also referred to as ‘consciousness’ in medical settings.

The nature of visual consciousness is yet to be fully elucidated and the current theories differ on the neural processes that underlie visual consciousness. According to the global neuronal workspace theory, the broadcasting and amplification of stimulus-specific information, specifically in prefronto-parietal areas, is what allows a visual stimulus to enter consciousness (Dehaene & Changeux, 2011; Salti, Monto, Charles, King, Parkkonen, & Dehaene, 2015). In contrast, the higher-order theory of consciousness asserts that visual consciousness does not involve the amplification or broadcasting of stimulus-specific information (Lau & Rosenthal, 2011; also see Salti et al, 2015 for discussion). Rather, non-stimulus-specific information is added, marking the stimulus as ready to enter consciousness. While the global neuronal workspace and higher-order theories differ on the nature of visual consciousness, they agree that consciousness emerges at a late stage of processing.

Visual consciousness has been studied by examining correlates of consciousness in brain activity (e.g., Pitts, Metzler, & Hillyard, 2014; Lamy, Salti, & Bar-Haim, 2009). In humans, this research most often has taken a univariate approach, examining regional brain activity measured with fMRI (cf. Haynes, 2009). Using this approach, for example, activation in the lateral occipital complex (LOC) measured using fMRI has been linked to visual consciousness (Grill-Spector, Hendler, Kushnir, & Malach, 2000). In EEG, a positive component called the P3b, has been found to occur when visual consciousness is present (Dehaene & Changeux, 2011; Lamy et al, 2009). The P3b component emerges at 300-500 ms post-stimulus onset, indicating that visual consciousness is likely to arise during this late time window. Yet, an earlier component has also been linked to visual consciousness (Pitts et al, 2014). This component, coined the visual awareness negativity, emerges at 200-240 ms post-stimulus onset, and has been found to correlate with consciousness regardless of the task relevance of the stimuli.

Perceptual and cognitive phenomena, such as visual consciousness, may not be characterised by any one single activation, but by the pattern of multiple activations across the brain (Haynes, 2009). This idea lends itself to multivariate pattern analysis (MVPA) or “decoding” approaches, which study distributed patterns of activity in the brain. In fMRI studies, decoding studies have shown that it is possible to predict various types of stimulus-specific information such as stimulus category and stimulus location from brain activity (e.g., Carlson, Schrater, & He, 2003; Cox & Savoy, 2003; Haxby, Gobbini, Furey, Ishai, Schouten, & Pietrini, 2001; Haynes, 2015; Kamitani & Tong, 2005; Kriegeskorte, Goebel, & Bandettini, 2006; Shinkareva, Malave, Just, & Mitchell, 2012). Moreover, the decoding performance has been shown to change as a function of consciousness (e.g., Williams, Dang, & Kanwisher, 2007; Bode, Bogler, Soon, & Haynes, 2012). Williams et al., for example, found that patterns of activity in the Lateral Occipital Complex (LOC) could reliably predict object category only when the participants were consciously aware of the stimuli. Activity in V1, in contrast, could be used to predict object category regardless of whether the stimulus was consciously perceived. Similarly, Bode et al., (2012) found that LOC activity could predict stimulus category only when visual consciousness was present. Studies using decoding methods thus corroborate what has been previously reported by univariate studies, in particular that the LOC is implicated in the conscious perception of objects.

To date, there has been relatively little research using decoding methods to investigate the temporal dynamics of visual consciousness. Decoding methods provide a means to study the dynamics of visual consciousness processing by revealing what information is being represented by the brain, and also when. For example, one study showed that stimulus information can be decoded more than 1000 ms after stimulus onset, both when stimuli are consciously perceived and when they are not (King, Pescetelli, & Dehaene, 2016). Further, the time that decoding performance between consciously perceived and non-consciously perceived stimuli diverges can be used as an indicator of the time visual consciousness emerges. Using this approach, Salti et al., (2015) found that visual consciousness for object location emerges at 270 ms post-stimulus onset. Their findings thus suggest that visual consciousness for object location, typified as greater decoding performance for consciously perceived stimuli, emerges around 270 ms post-stimulus onset.

The present study aimed to investigate the time that visual awareness of stimulus category emerges. We recorded magneto-encephalography (MEG) data while participants completed a visual categorisation task. We manipulated participants’ consciousness of the stimuli using a standard backward masking paradigm (Breitmeyer & Ögmen, 2006). Consciousness was measured using three different methods; by an objective measure (behavioural categorisation accuracy), by a subjective measure (self-report of visibility), and by a combination of both the objective and subjective measures. These three different methods were used to address concerns that forced-choice categorisation alone is not an adequate measure of visual consciousness (Dehaene & Changeux, 2011). Using decoding to analyse the MEG data, we identified the time at which the neural signal started to differ between trials where visual consciousness was present, and trials where visual consciousness was not evident. We found that visual consciousness is characterised by an increased decodability of stimulus information, emerging around 180 – 230 ms post-stimulus onset.

## 2. Methods

The aim of the current study was to disentangle conscious from unconscious processing in visual object categorisation using backward masking paradigm. Participants performed a three-way categorisation task for three artificial categories of objects (Spikies, Smoothies, Cubies (Op de Beeck, Baker, DiCarlo, & Kanwisher, 2006)). The experimental session consisted of two phases. In the first phase, participants were familiarised with the task while we adaptively estimated their individual contrast threshold so that their accuracy was maintained at 50%. Then, in the second phase, participants performed the categorisation task while we recorded their brain activity in response to the stimuli with magnetoencephalography (MEG).

### 2.1 Participants

Eight healthy adults (5 female) participated in the study. All were between the age of 18 and 24 (mean age = 20.38 years, *SD* = 1.77 years). Participants were fluent in English and had normal or corrected-to-normal vision. One participant was left-handed. The participants gave informed consent in writing prior to their participation and were financially reimbursed for their time. The study was conducted with the approval of the Human Research Ethics Committee at the University of Sydney and Macquarie University.

### 2.2 Stimuli

The stimuli were novel objects belonging to one of three categories: cubies, smoothies, and spikies (Figure 1A), artificially generated using Matlab (Op de Beeck et al., 2006). There were 210 visually different exemplars in each object category (Figure 1A). Stimuli were presented in greyscale on a black background (Figure 1B), at the centre of the screen (visual angle: 3.23°×2.87°). The stimuli were masked using greyscale random dot masks, constructed by assigning random values to the pixels of a 100×100 image, displayed at the same size and location as the stimuli. The same stimuli and masks were used in the familiarisation and test phase of the study. During the familiarisation phase, the contrast in which the stimuli were presented was calibrated for each participant, so that they would correctly categorise the object 50% of the time (chance level = 33%). During the test phase, the contrast also varied from trial to trial, to ensure that the participants would correctly categorise the object approximately 60% of the time using QUEST (Watson & Pelli, 1983). We used 50% in the familiarisation phase to make the training engaging and challenging, and then increased the threshold 60% to get a good distribution of correct and incorrect trials for the main experiment. The experiment was run in Matlab R2011b, using the Psychophysics Toolbox version 3.0.10 (Brainard, 1997; Kleiner et al., 2007; Pelli, 1997). The stimuli were projected onto a screen inside the magnetically shielded room using an EPSON EB-G7400U projector. The participants reported their responses using a 4-button cylinder box.

**Figure 1.**
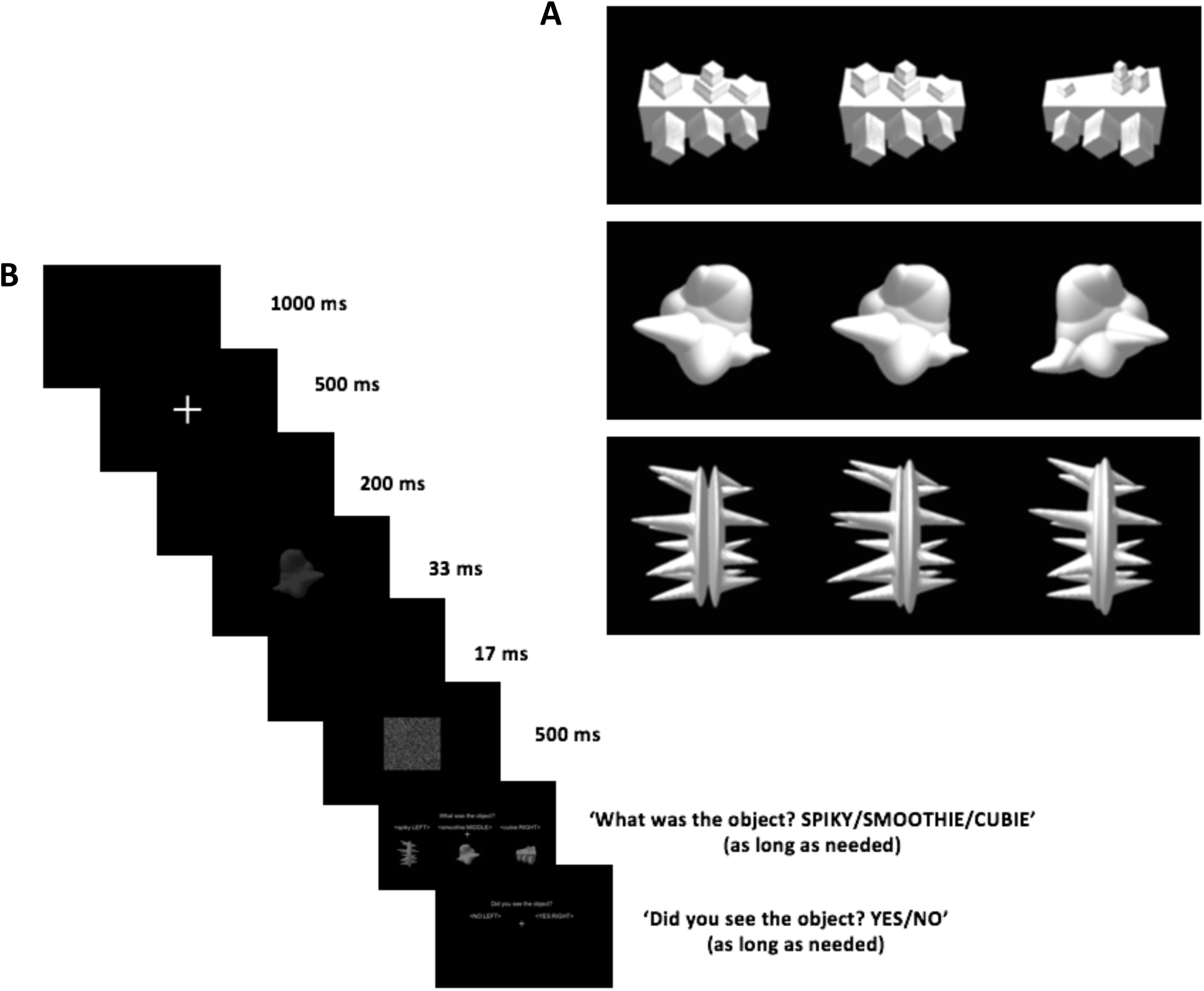
A. The three object categories: cubie, smoothie, and spiky. For each category, there were 210 visually different exemplars. Here we show three examples of each category. B. The experiment paradigm in the test phase. On each trial, participants were shown an object, followed by a mask. Participants were instructed to report the category of the object (response mapping was randomised between blocks), and finally they reported whether they perceived the stimulus or not.

### 2.3 Procedure

At the start of the experiment, the participants were fitted with a cap containing 5 marker coils to monitor head movement. Their head shape was also digitised using the Locator programme with Fastrak Polhemus (version 5.5.2) to check the location and alignment of the head in the scanner. The participants laid in a supine position in the MEG scanner, in a dimly lit magnetically shielded room (Fujihara Co. Ltd., Tokyo, Japan). The participants were instructed to minimise their head movements whilst inside the scanner. The recordings were made using a whole-head MEG system containing 160 axial gradiometers (Model PQ1160R-N2, KIT, Kanazawa, Japan). MEG signal was continuously sampled at 1000 Hz, band-pass-filtered online between 0.03Hz and 200 Hz.

The experiment employed a backward masking paradigm. The target stimulus was presented for a brief duration followed by a mask at the same location where the stimulus was previously displayed (Figure 1B).

The first part of the experimental session was a familiarisation phase. During this phase, there was a 1000 ms interval at the start of each trial, followed by a fixation cross presented at the centre of the screen for 500 ms. This was followed by another 200 ms interval with a blank screen. Subsequently, the target stimulus was presented for 33 ms. After target stimulus offset, there was a delay of 17 ms before a random-dot mask appeared. The mask was presented for 500 ms, at the same location where the target stimulus previously appeared. Following mask offset, participants were prompted to categorise the object shown in the trial. Participants were allowed as much time as needed to categorise the object. Once the participants had entered a response, the next trial started.

The familiarisation phase consisted of 100 trials, 25 of which were control trials, which did not contain a stimulus. In these trials, presentation of the target stimulus was replaced with a blank screen, which lasted for 33 ms (the same duration as stimulus presentation in target trials). The remaining 75 trials were target trials, where the target stimuli were presented. In these trials, the stimulus was either a spiky, smoothie or cubie. All three categories were presented equally often, to eliminate bias for any particular category. Stimuli for each category were randomly drawn from the stimuli pool described in Section 2.2. All four types of trials (spiky, smoothie, cubie, and control trials) were presented in a random order during the familiarisation phase. The familiarisation phase lasted approximately half an hour. There was no MEG acquisition during this phase.

Upon completion of the familiarisation phase, the participants commenced the test phase. The test phase followed a similar procedure as the familiarisation phase, except that after the categorisation question, the participants were also asked: “Did you see the object?”. The participants selected either the “Yes” or “No” response. They were instructed to respond “Yes” only when they had seen the stimulus and were also able to identify what category it was. As with the categorisation question, participants were allowed as much time as necessary to respond to this question.

The test phase consisted of seven blocks. Each block lasted for approximately 8 minutes and was comprised of 168 trials (42 spiky trials, 42 cubie trials, 42 smoothie trials and 42 control trials). At the start of each block, the response mapping for the categorisation question was changed to ensure that motor response could not act as a confound. The response mapping was changed after every block in a random order. The test phase lasted an hour including the breaks between blocks.

### 2.4 Analysis

#### 2.4.1 Pre-processing

At the time of the experiment, 9 MEG channels were undergoing maintenance and the analysis was performed on the remaining 151 channels. The data were down-sampled to 100Hz (10ms resolution). Stimulus onset times were determined using a photodiode located in the corner of the display in the magnetically shielded room. MEG recordings were sliced into epochs starting from 100 ms prior to stimulus onset and ending at 800 ms post-stimulus onset. Pre-processing was performed in Matlab R2017, using the FieldTrip Toolbox (version 20170502) (Oostenveld, Fries, Maris, & Schoffelen, 2011). No further preprocessing steps were applied to the data.

#### 2.4.2 Decoding

We performed a time-series decoding analysis on the preprocessed data (Grootswagers, Wardle, & Carlson, 2017), implemented in CoSMoMVPA (Oosterhof, Connolly, & Haxby, 2016). After discarding control trials, we decoded the category of the stimulus for each participant over the time course of the trial. We used linear discriminant analysis (LDA) classifiers as implemented in CoSMoMVPA. The classifier was trained at every time point in the epoch, using the activation values from all MEG channels. The decoding performance was examined using a leave-one-block-out cross-validation method, training the classifier on all-but-one blocks, testing it on the remaining block, and repeating this leaving every block out for testing once. We applied this analysis on all pairwise combinations of category pairs (i.e., spiky versus smoothie, spiky versus cubie, and smoothie versus cubie) and report the mean cross-validated decoding performance across pairwise combinations.

Stimuli were presented at a varying contrast throughout the experiment (using the QUEST adaptive procedure). We therefore took the following steps to control for contrast: firstly, we excluded the first block of each participant, where the QUEST procedure had not yet converged, and contrast was more variable. Secondly, we exactly matched the contrast of correct and incorrect trials for the analysis; for each trial with an incorrect response, we selected a correct response trial that was presented at the exact same contrast value. If no matching trial was found, the trial was excluded. On average, this procedure retained 74.82% of trials (mean±*SD*: 377.08±78.84 trials). This approach ensured that the decoding procedure was performed not only on equal numbers of correct and incorrect trials (thus avoiding classifier bias), but also that the correct and incorrect trials had the exact same contrast values and distributions. Within the cross-validation procedure, the classifier was trained on all remaining trials. To examine the difference between conscious and unconscious processing, we grouped the trials in the test set and assessed their decoding performance separately according to the following three comparisons:

1. ‘correct’ versus ‘incorrect’ trials (objective measure)
2. ‘seen’ versus ‘unseen’ trials (subjective measure)
3. ‘correct-seen’ versus ‘incorrect-unseen’ trials (combined measure)

#### 2.4.3 Statistical testing

At each time point in the response, we tested whether decoding accuracy was at chance-level (H0), or above chance (H1). We also tested whether the decoding performance between groupings (e.g., correct versus incorrect) was the same (H0) or different (H1). To compare hypotheses, we used Bayes Factors (BF), which quantify the evidence for one hypothesis over the other (Jeffreys, 1998; Morey, & Rouder, 2011; Rouder, Speckman, Sun, Morey, & Iverson, 2009; Wagenmakers, 2007; Wetzels et al., 2011). In the Bayesian framework, a BF of 3 indicates H1 is three times more likely than H0, and a BF of 1/3 indicates the opposite. A BF>3 or BF<1/3 is generally considered as substantial evidence (roughly comparable to a p-value < 0.01), and BF>10 or BF<1/10 as strong evidence (roughly comparable to a p-value < 0.001) for H1 or H0, respectively (Dienes, 2016; Jeffreys, 1998; Wagenmakers, 2007; Wetzels et al., 2011). Note that the Bayes factors are continuous degrees of evidence, and the two levels of thresholding are mainly used for visualisation purposes. We did not treat these thresholds as hypothesis testing at the singe time point level, and instead consider the evidence across multiple time points. This means that isolated time points that reach the threshold are not treated as evidence for a hypothesis if the evidence in the surrounding time points goes in the opposite direction.

We constructed a uniform prior for H1 with an upper bound set at 100% in the case of decoding accuracy, and at 50% for the difference between accuracies (Dienes, 2008; 2014). Instead of using chance as lower bound for H1, we constructed a conservative estimate of the lower bound using a permutation test (Maris & Ooostenveld, 2007; Stelzer, Chen, & Turner, 2013) as follows: for each participant, we created 100 null-results by performing the classification analysis on shuffled class labels. We then sampled at random 5,000 times from the individual participant null-distributions and computed the mean decoding performance, resulting in a group level null-distribution (Maris & Oostenveld, 2007). We used the group-level decoding accuracy at the 95^th^ percentile of this null-distribution as the lower bound of the prior for H1. When comparing the difference in decoding performance between groupings (e.g., correct versus incorrect), we created in a similar way a group-level null-distribution of differences and used the 95^th^ percentile of this distribution as lower bound for the difference between accuracies.

#### 2.4.4 Exploratory analysis

To explore the source of the decodable signal, we performed a channel-space searchlight analysis for the the combination of both the objective and subjective measures (i.e., correct seen versus incorrect not seen). For a given channel, we took the 4 closest neighbouring channels and performed the same decoding procedure on this local cluster of channels. The decoding accuracy was then stored at the centre channel. This process was repeated for all channels, yielding a scalp map of decoding accuracies for every time point.

## 3. Results

The aim of the study was to investigate the temporal dynamics of visual consciousness. We operationalised visual consciousness using three different methods: (1) objective measure alone (i.e. categorisation accuracy), (2) subjective measure alone (i.e. self-report of visibility); (3) the combination of both the objective and subjective measures. We decoded the stimulus category (spikey, smoothie, cubie) and then compared the decoding performance in three sets of comparisons corresponding to the three operationalised definitions of consciousness: (1) between trials where participants responded correctly in the categorisation task and those where they responded incorrectly (‘correct’ vs ‘incorrect’ trials; Figure 2A); (2) between trials where participants reported having seen the stimulus and those where they reported not having seen it (‘seen’ vs ‘unseen’; Figure 2B); (3) between trials where participants responded correctly in the categorisation task and also said they saw the stimulus and trials where participants neither responded correctly in the categorisation task nor reported seeing the stimulus (‘correct-seen’ vs ‘incorrect-unseen’; Figure 2C).

**Figure 2.**
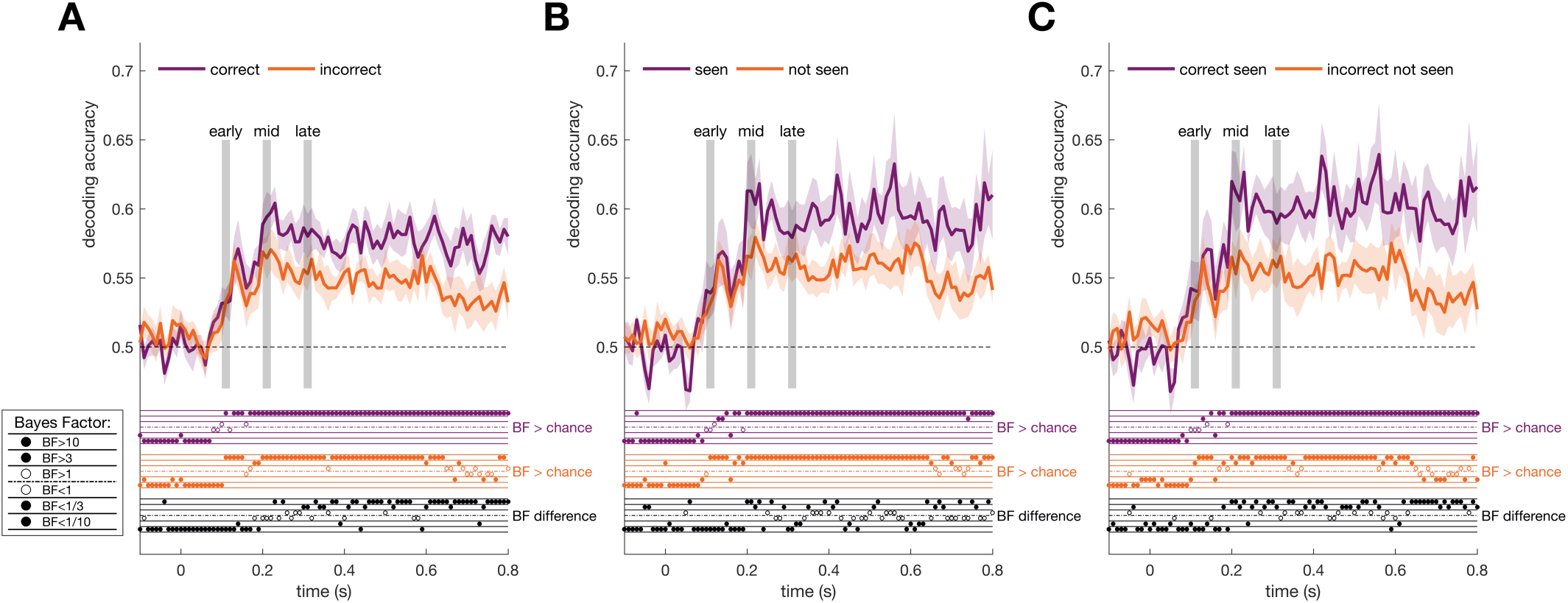
The time course of decoding performance for: A. ‘correct’ vs ‘incorrect’ trials; B. ‘seen’ vs ‘unseen’ trials; C. ‘correct-seen’ vs ‘incorrect-unseen’ trials. Shaded regions show ± 1 SEM across participants. The broken horizontal line indicates chance level. The y-axis indicates decoding performance, with 1 being 100% accuracy, and 0.5 indicating 50% accuracy. The x-axis indicates the time course of the trials in seconds relative to stimulus onset. Bayes Factors (BF) are indicated by the dots above the x-axis of each graph. BF were thresholded at 1/10, 1/3, 1, 3, and 10 (see inset). A BF of 1/3 or below indicates evidence for the null hypothesis (filled dots in the bottom two rows), and a BF of 3 or above indicates evidence for the alternative hypothesis (filled dots in the top two rows), and BF between those values reflects insufficient evidence for either hypothesis (open dots in the two middle rows). Purple and orange dots in each graph indicate the BF for above-chance decoding for the purple and orange lines in that graph, respectively. Black dots indicate the BF for the difference in decoding performance between the purple and orange conditions in that graph. The shaded vertical grey areas show the three time points shown in Figure 3 for the exploratory channel-searchlights.

### Objective measure

In the first comparison (Figure 2A), visual consciousness was operationalised by the objective measure: the participants’ accuracy in the categorisation task. Trials where participants responded correctly (‘correct’ trials) showed decoding performance that was above chance starting from 110 ms post-stimulus onset (BF= 10.52). Trials where participants responded incorrectly (‘incorrect’ trials) also showed above chance decoding performance starting from 110 ms post stimulus onset (BF= 12.16). The ‘correct’ trials were first observed to have higher decoding performance than the ‘incorrect’ trials at 230 ms post-stimulus onset (BF = 47.00). Between 230 ms and 410 ms post-stimulus onset, this difference was inconsistent, but from 410 ms onwards, the ‘correct’ trials consistently had better decoding performance compared to the ‘incorrect’ trials. Prior to 190 ms, there was evidence for the null hypothesis or no difference between the ‘correct’ and ‘incorrect’ trials (BF < 1/3).

### Subjective measure

In the second comparison (Figure 2B), visual consciousness was operationalised by the subjective measure: participants’ subjective report of visibility. In trials where participants reported that they saw the stimulus (‘seen’ trials), the decoding performance rose above chance from 130 ms post-stimulus onset (BF = 5.94). In ‘unseen’ trials, the decoding performance was above chance from 110 ms post-stimulus onset (BF = 17.77). Decoding performance for ‘seen’ trials was better than that for ‘unseen’ trials, with this difference emerging at 200 ms post-stimulus onset (BF = 14.65). However, this difference was not as consistent throughout the rest time series as it was for the first comparison.

### Combined measure

In the third comparison (Figure 2C), visual consciousness was operationalised by a combination of both the objective and subjective measures: categorisation accuracy and self-report of visibility. In the ‘correct-seen’ trials, the decoding performance was above chance from 130 ms post-stimulus onset (BF = 3.54). The ‘incorrect-unseen’ trials showed decoding performance above chance starting from 110 ms post-stimulus onset (BF = 5.60). From 180 ms, there was a difference in decoding performance with the ‘correct-seen’ trials showing better decoding performance than ‘incorrect-unseen’ trials from this time onwards (BF = 9.45). There was also evidence for no difference between ‘correct-seen’ and ‘incorrect-unseen’ trials prior to 170 ms.

### The Neural Source of Decodable information: An exploratory analysis

An exploratory analysis using channel-searchlights (Figure 3) indicated that during the middle time period (200-220 ms), decodable stimulus information was found around occipital channels in the ‘incorrect-unseen’ trials, and from frontal and occipital channels in the ‘correct-seen’ trials. Compared to the ‘incorrect-unseen’ trials, there seemed to be more decodable stimulus information coming from frontal channels. During the late time period (300-320 ms), decodable stimulus information was found in the occipital and frontal channels in both the ‘correct-seen’ and ‘incorrect-unseen’ trials, and there was a greater amount of decodable information from the occipital channels in the ‘correct-seen’ trials.

**Figure 3:**
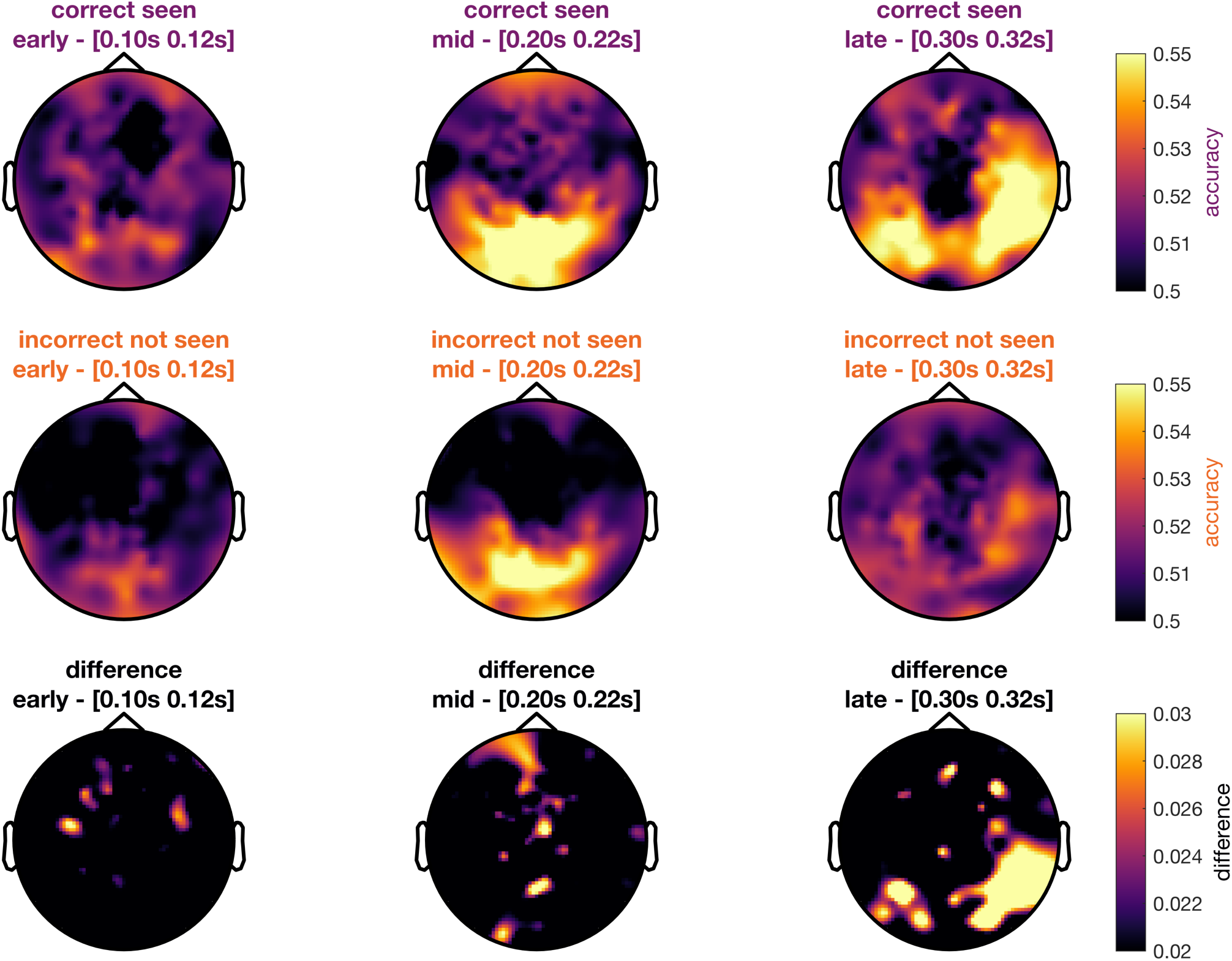
Result for the exploratory searchlight analysis for ‘correct-seen’ versus ‘incorrect-unseen’ comparison. These maps show channel decoding accuracies for the three timepoints annotated in Figure 2. The first row shows decoding accuracy for ‘correct-seen’ trials, the second row for ‘incorrect-unseen’ trials, and the bottom row shows the difference.

## 4. Discussion

This study investigated the information represented by the brain during conscious and unconscious processing of visual objects. In a MEG recording session, we showed participants stimuli at threshold, such that only on a subset of trials the stimuli reached conscious awareness, and had participants give objective (i.e., categorisation accuracy) and subjective (i.e., self-report of visibility) reports on the stimulus they were viewing. In our analysis, we then operationalised consciousness using objective, subjective, and a combined objective-subjective measure to study how stimulus information was represented in the brain during consciousness and unconscious processing.

### 4.1. Visual consciousness characterised by increased decodability for stimulus information

Across all the definitions of consciousness, we found consistent patterns of results regarding the information represented during conscious and unconscious processing. Irrespective of definition, we could decode object category information from both conscious and unconscious trials. Notably, showing that we can decode stimulus information during unconscious trials demonstrates that the brain represents object information even if the stimulus does not reach conscious awareness. When consciousness was operationalised by the objective measure (categorisation accuracy), we found that decoding performance for correct trials was higher than ‘incorrect’ trials starting from 230 ms post-stimulus onset. A similar pattern of results emerged for the subjective and combined objective-subjective definitions of consciousness. In both cases, we observed higher decoding performance for “conscious” than “unconscious” trials. Collectively, these findings indicate that the difference between conscious and unconscious processing is better characterised as a difference in the strength of the stimulus representation, which is that information is enhanced (i.e., more decodable) during conscious processing.

### 4.2 Stimulus-related information is processed by the brain with conscious awareness of the stimulus

Stimulus information was present when visual consciousness was considered absent using all three operationalised definitions, indicating that some processing is completed by the brain independent of visual consciousness. These results corroborate fMRI decoding studies showing stimulus information is represented in the brain even when the stimulus is not consciously accessible. Williams et al., (2007), for example used an objective measure of consciousness (i.e., behavioural performance) to show that object category information could be decoded from primary visual cortex even when subjects incorrectly reported the stimulus category. Our study further showed that when consciousness was operationalised using subjective report (i.e., seen/unseen trials), stimulus information was decodable during unconscious processing. These results echo the findings of King et al., (2016), who showed that stimulus information is encoded and maintained in the brain up to 1150 ms post-stimulus onset, irrespective of the subjective reports. Finally, we also found that stimulus information was decodable for unconscious trials using the combined objective-subjective measure (i.e., ‘incorrect-unseen’ trials). Collectively, our findings show that irrespective of the method used to operationalise visual consciousness, stimulus information is represented by the brain even when the stimulus is not consciously accessible to the observer.

### 4.3. Visual consciousness emerges between 180 – 230 ms post-stimulus onset

Conscious trials showed higher decoding performance regardless of the operationalised definition of consciousness, a difference that emerged between 180 – 230 ms post-stimulus onset. This time is notably earlier than the 270ms estimate reported in a decoding study by Salti et al., (2015). There are several possible explanations for this discrepancy. Firstly, Salti et al. displayed their stimuli in the periphery, whereas in the present study stimuli were displayed at the fovea. Visual acuity is lower in the periphery (Anstis, 1974; 1998), thus one explanation is the peripheral stimuli used by Salti et al. were weakly represented and/or took longer to be processed. Due to the reduced fidelity in processing stimuli in the periphery, visual consciousness thus might have been found to emerge at a later time.

Secondly, Salti et al. (2015) divided the time course into (four) discrete time windows, while the present study investigated time series by measuring decoding accuracy at each time point. Notably, the time for visual consciousness in our results was 180 – 230 ms, which falls at the mid-point of the third time window defined by Salti et al. (i.e., 162-271 ms post-stimulus onset). The second time window used by Salti et al. potentially could have had added noise, which rendered the difference between conscious and unconscious processing insignificant. Thus, a second explanation is that our fine-grained temporal resolution led to finding differences at an earlier time.

Finally, the two studies examined different stimulus properties. In Salti et al., the stimulus property of interest was stimulus location, whereas we investigated stimulus category. The discrepancy in the findings might be explained by the fact that consciousness for category emerges at an earlier time than that for stimulus location. This possibility contradicts earlier findings showing that the decoding onset for stimulus category emerges after that for stimulus location (Carlson, Hogendoorn, Kanai, Mesik, & Turret, 2011). Moreover, it is generally accepted that stimulus location is represented early (i.e., primary visual cortex), while object category information is represented at a later stage in the visual hierarchy (lateral occipital cortex and inferior temporal cortex). The explanation that the conscious representation of location precedes the representation of category thus contradicts both previous decoding studies and accepted knowledge of the visual hierarchy. We, therefore, view this latter explanation as possible, but not plausible. Nevertheless, future work could investigate these three possible explanations for the difference between Salti et al. and our study’s findings.

Other studies have taken a univariate analysis approach with EEG to study the brain dynamics of consciousness. Dehaene and Changeux (2011) and Lamy et al. (2009), for example, reported the P3b correlated visual consciousness. The P3b is an event-related potential (ERP) with onset between 300-500 ms. This timing is notably later than the time window reported in this study (between 180-230 ms). In contrast, Pitts, Metzler, et al. (2014) and Pitts, Padwal, Fennelly, Martínez, and Hillyard (2014) showed that the visual awareness negativity (VAN) correlated to visual consciousness. The onset of the VAN is approximately 200ms, which coincides more closely with our estimate of the time of the emergence for visual consciousness.

### 4.4. Stimulus information associated with visual consciousness does not preclude the existence of a non-stimulus specific ‘tag’ for consciousness

The difference between the presence and absence of visual consciousness manifested in the strength of decoding performance. Visual consciousness thus correlates with increased decoding performance. This observation, to some extent, supports the global neuronal workspace theory, which proposes that visual consciousness emerges due to the amplification of stimulus-specific information (Dehaene & Changeaux, 2011). Our exploratory channel-searchlight results indicated that during the middle time period, this amplification of neural information found had its source in the frontal lobe. However, as the searchlight analysis were exploratory in nature, it was not known whether the neural information found was strongly related to the prefrontal cortex, which is implicated in the global neuronal workspace theory. Moreover, the finding that visual consciousness relates to the strength of the representation does not preclude the possibility that additional non-stimulus-specific signals are also involved, as proposed by the higher order theory (Lau & Rosenthal, 2011; see Salti et al., 2015 for discussion). Such non-stimulus-specific signals, could play a dual role by ‘tagging’ certain stimuli as ready for conscious perception, and simultaneously contributing to the amplification of stimulus-specific information.

### 4.5. The contribution of attention, memory and decision-making

A limitation in the present study is that it did not isolate the contribution of attention, memory and decision-making to the results. All these factors often co-occur with visual consciousness, yet are not visual consciousness per se (Aru, Bachmann, Singer, & Melloni, 2012; de Graf, Hsieh, & Sack, 2012; Lamme, 2006; Tallon-Baudry, 2012). Attention, in particular, has been shown to enhance neural activity in response to stimulus categories (Desimone & Duncan, 1995; Kastner & Ungerleider, 2000; O’Craven, Downing & Kanwisher, 1999). Moreover, memory is often required to maintain the conscious percept for subsequent reporting, and quite often the reporting process involves an explicit decision made by the participants. As a result, these additional factors also could mediate the observed relationship between visual consciousness and neural activity. It is therefore difficult to disentangle whether the difference in neural activity between ‘conscious’ and ‘non-conscious’ conditions is due to visual consciousness alone, or other concomitant factors such as attention, memory, and decision-making.

### 4.6. Conclusion

The present study aimed to examine the dynamics of visual consciousness by studying the brain’s representation of conscious and unconscious stimuli. Across three different methods of operationalising visual consciousness, we found that conscious awareness is characterised by increased decodability of neural signals encoding stimulus information. We found that this difference between conscious and unconscious processing emerges between 180 – 230 ms post-stimulus onset. Given that factors such as attention, memory and decision-making may have contributed to the findings, care must be taken when attributing the observed findings to visual consciousness alone. Nonetheless, our results corroborate existing literature on the neural characteristics of visual consciousness, and provide new evidence that visual consciousness may emerge earlier than previously established.

## Acknowledgements

This research was supported by an Australian Research Council Future Fellowship (FT120100816) and an Australian Research Council Discovery project (DP160101300) awarded to T.A.C. The authors acknowledge the University of Sydney HPC service for providing High Performance Computing resources. The authors declare no competing financial interests.

